## Supplementary Figures for "A broad mutational target explains an evolutionary trend"

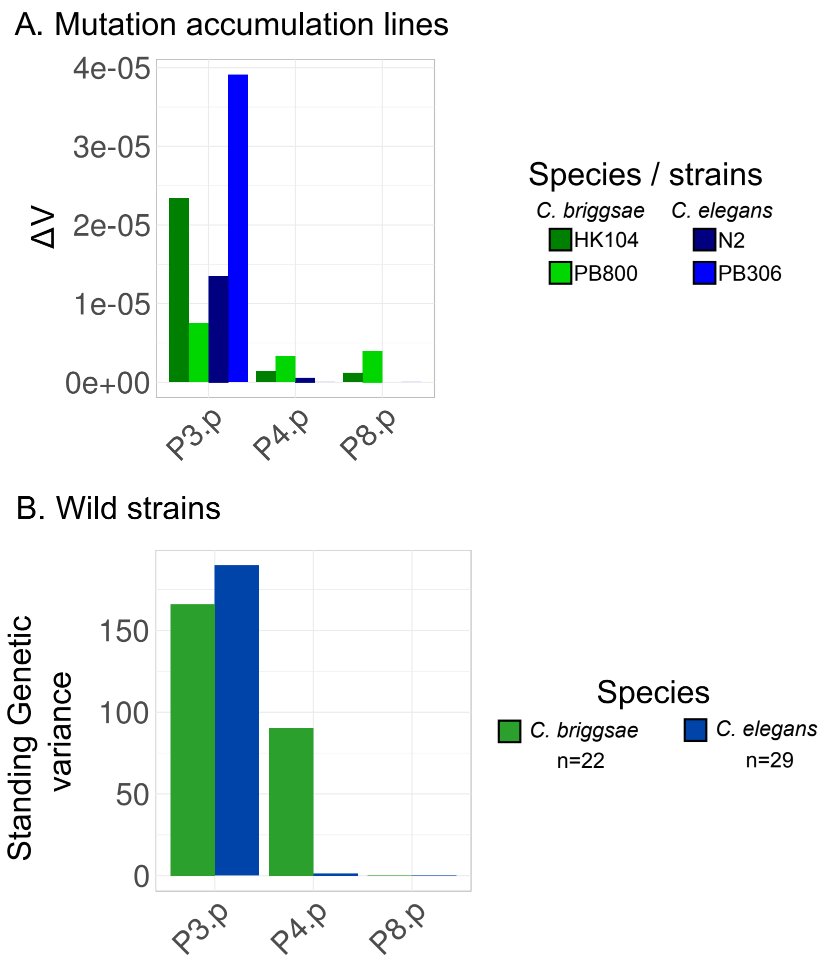
A. Mutation accumulation lines
ΔV
4e-05
3e-05
2e-05
1e-05
0e+00
P3.p
P4.p
P8.p
Species / strains
C. briggsae
C. elegans
HK104
PB800
N2
PB306
B. Wild strains
Standing Genetic variance
150
100
50
0
P3.p
P4.p
P8.p
Species
C. briggsae
n=22
C. elegans
n=29

Figure S1

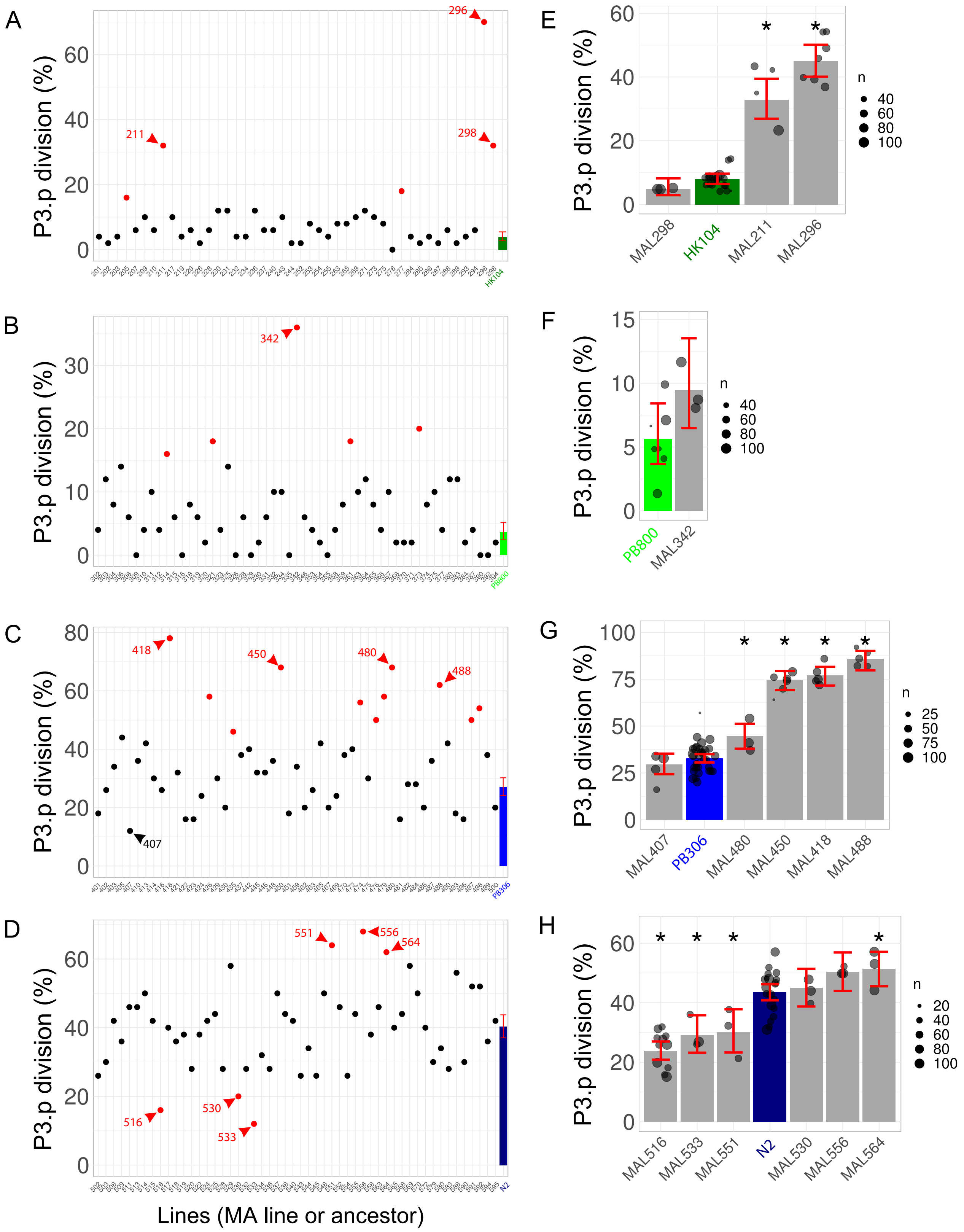
A
B
C
D
E
F
G
H
P3.p division (%)
Lines (MA line or ancestor)
296
211
298
342
418
450
480
488
407
551
556
564
516
530
533
MAL298
HK104
MAL211
MAL296
PB800
MAL342
MAL407
PB306
MAL480
MAL450
MAL418
MAL488
MAL516
MAL533
MAL551
N2
MAL530
MAL556
MAL564
n

Figure S2

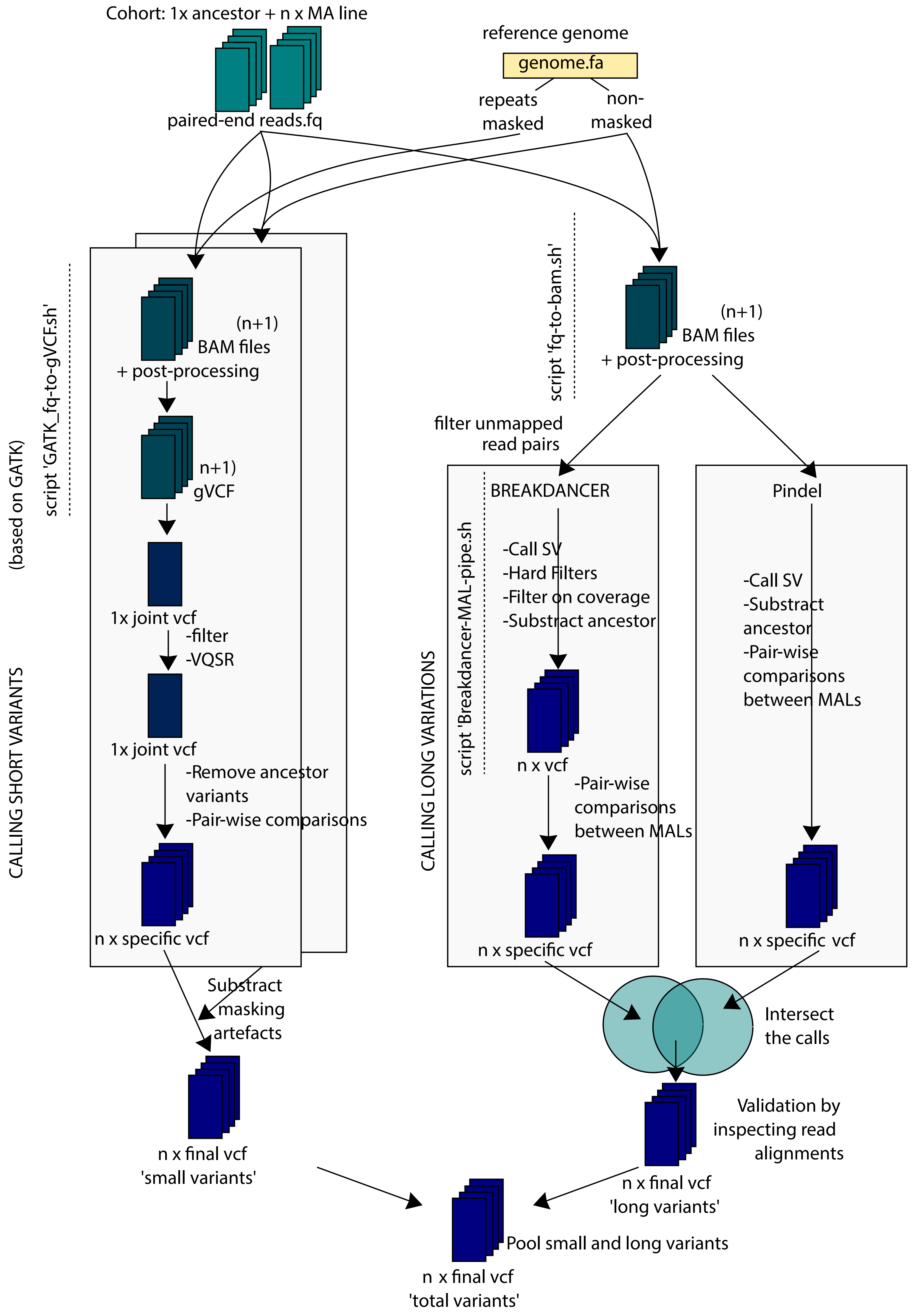
Cohort: 1x ancestor + n x MA line
paired-end reads.fq
reference genome
genome.fa
repeats masked
non-masked
script 'GATK_fq-to-gVCF.sh'
(based on GATK)
(n+1) BAM files + post-processing
n+1) gVCF
1x joint vcf
-filter
-VQSR
1x joint vcf
-Remove ancestor variants
-Pair-wise comparisons
n x specific vcf
CALLING SHORT VARIANTS
Substract masking artefacts
n x final vcf 'small variants'
script 'fq-to-bam.sh'
(n+1) BAM files + post-processing
filter unmapped read pairs
script 'Breakdancer-MAL-pipe.sh'
CALLING LONG VARIATIONS
BREAKDANCER
-Call SV
-Hard Filters
-Filter on coverage
-Substract ancestor
n x vcf
-Pair-wise comparisons between MALs
n x specific vcf
Pindel
-Call SV
-Substract ancestor
-Pair-wise comparisons between MALs
n x specific vcf
Intersect the calls
Validation by inspecting read alignments
n x final vcf 'long variants'
Pool small and long variants
n x final vcf 'total variants'

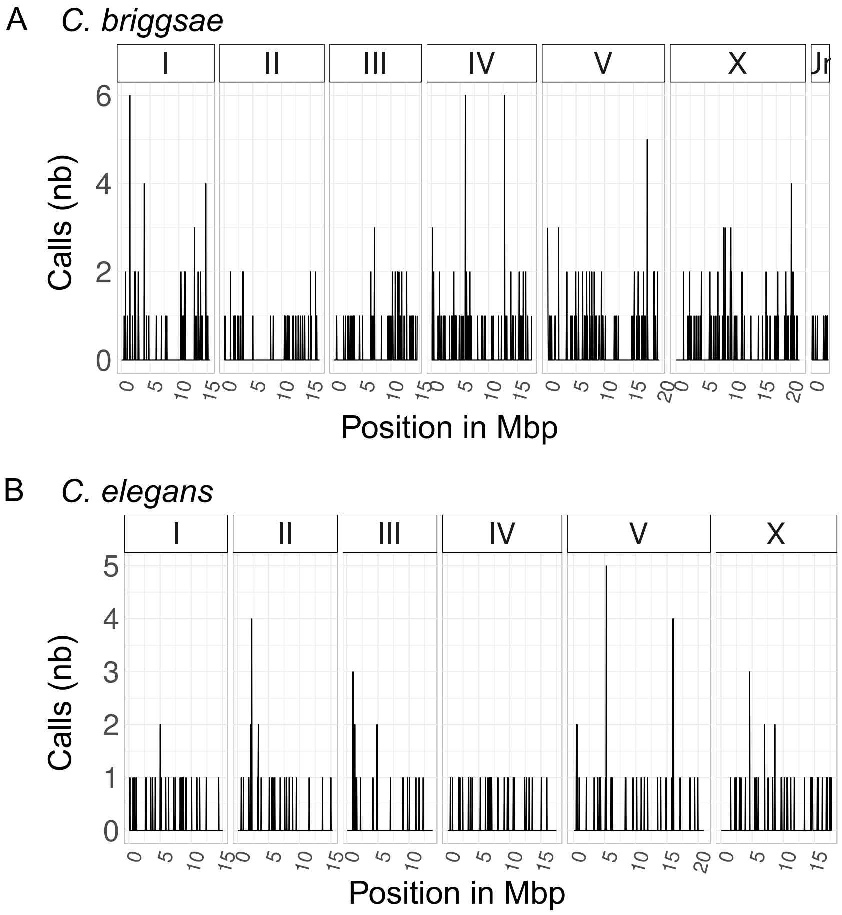
A
C. briggsae
I
II
III
IV
V
X
Calls (nb)
Position in Mbp
B
C. elegans
I
II
III
IV
V
X
Calls (nb)
Position in Mbp

Figure S4

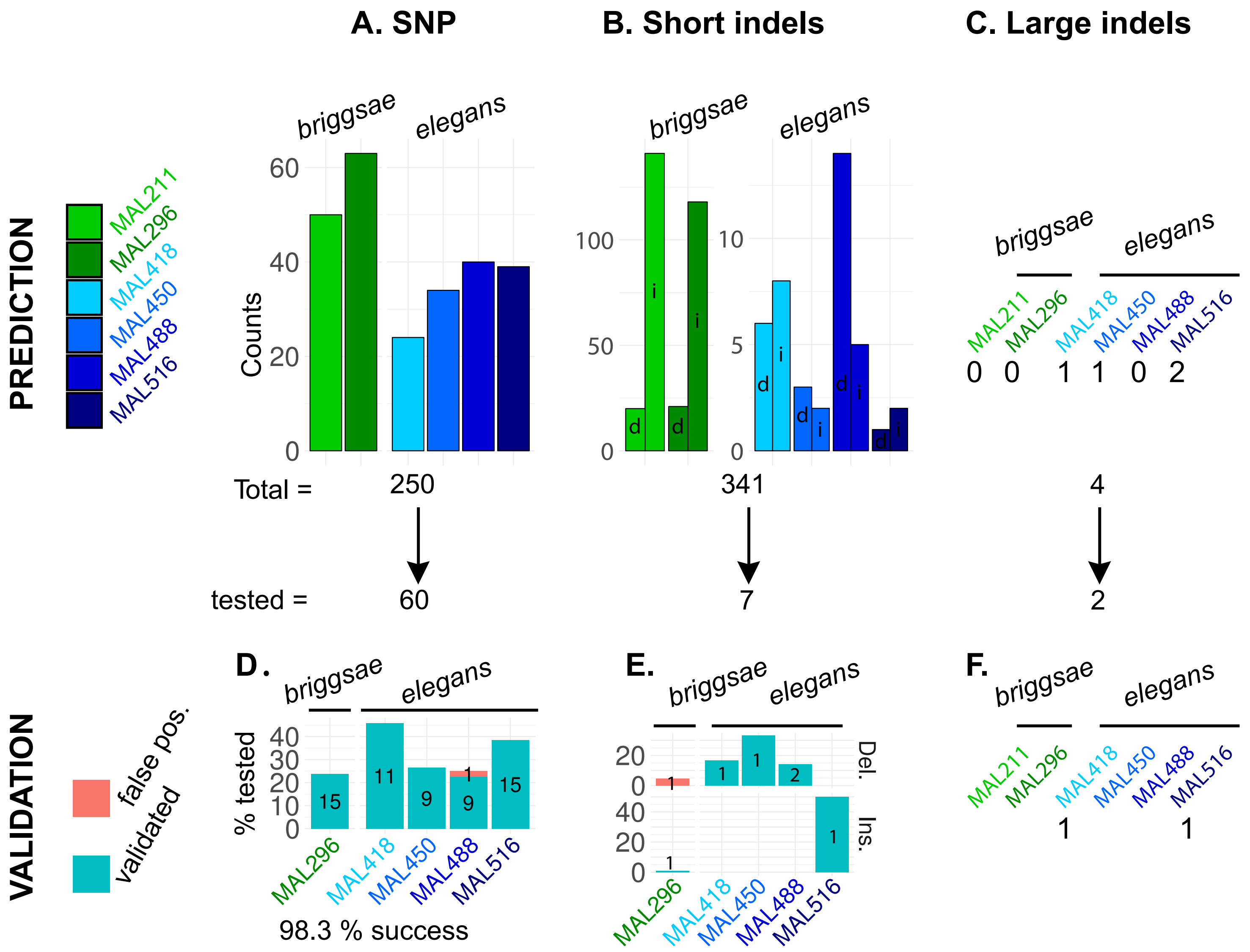

A. SNP
B. Short indels
C. Large indels
PREDICTION
MAL211
MAL296
MAL418
MAL450
MAL488
MAL516
briggsae
elegans
Counts
Total =
250
341
4
tested =
60
7
2
VALIDATION
false pos.
validated
D.
% tested
98.3 % success
E.
Del.
Ins.
F.


Figure S5

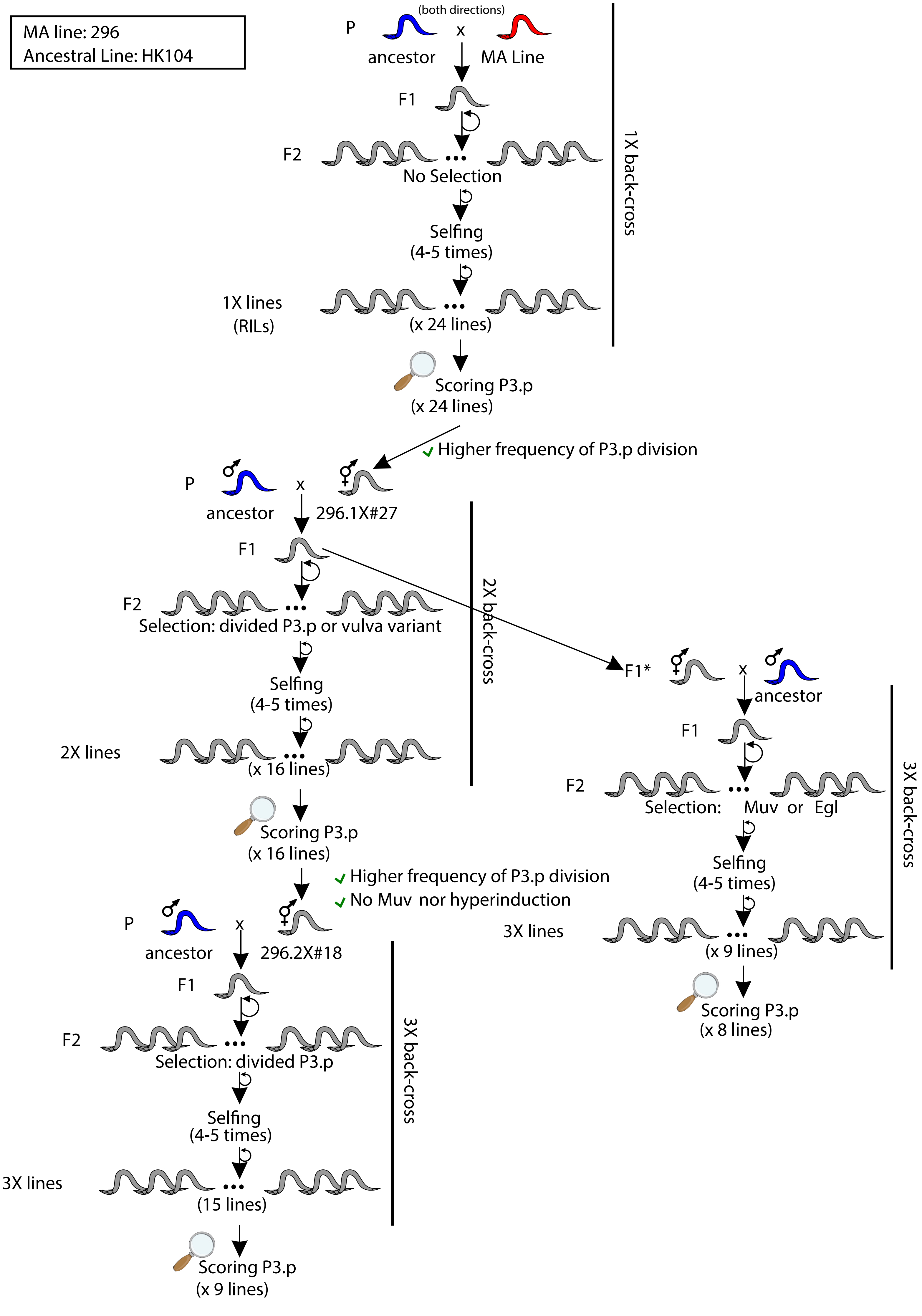

MA line: 296
Ancestral Line: HK104
(both directions)
P
x
ancestor
MA Line
F1
F2
No Selection
Selfing
(4-5 times)
1X lines
(RILs)
(x 24 lines)
1X back-cross
Scoring P3.p
(x 24 lines)
Higher frequency of P3.p division
P
x
ancestor
296.1X#27
F1
F2
Selection: divided P3.p or vulva variant
Selfing
(4-5 times)
2X lines
(x 16 lines)
2X back-cross
Scoring P3.p
(x 16 lines)
Higher frequency of P3.p division
No Muv nor hyperinduction
F1*
x
ancestor
F1
F2
Selection: Muv or Egl
Selfing
(4-5 times)
3X lines
(x 9 lines)
3X back-cross
Scoring P3.p
(x 8 lines)
P
x
ancestor
296.2X#18
F1
F2
Selection: divided P3.p
Selfing
(4-5 times)
3X lines
(15 lines)
3X back-cross
Scoring P3.p
(x 9 lines)

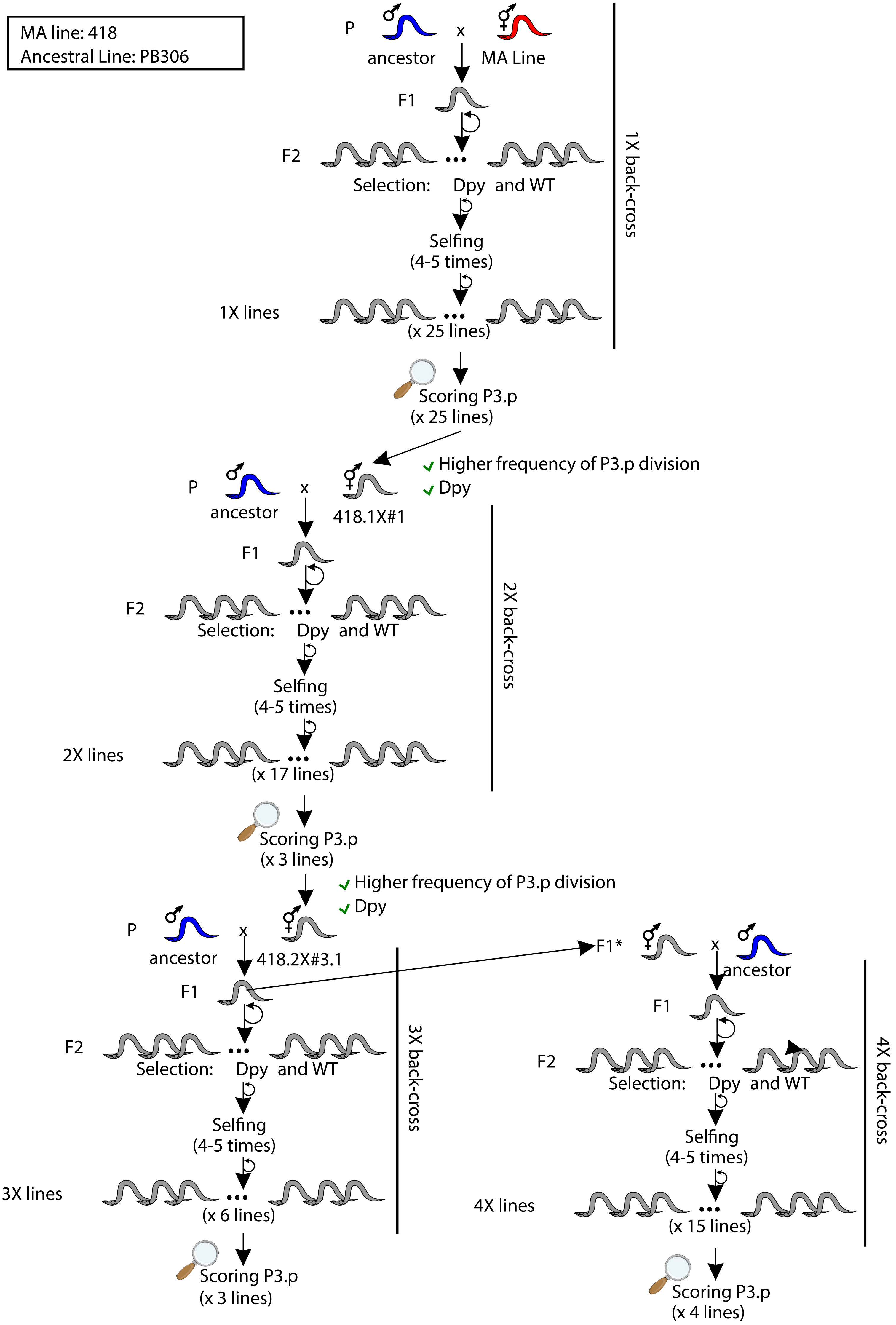
MA line: 418
Ancestral Line: PB306
P
ancestor
x
MA Line
F1
F2
Selection: Dpy and WT
Selfing
(4-5 times)
1X lines
(x 25 lines)
1X back-cross
Scoring P3.p
(x 25 lines)
Higher frequency of P3.p division
Dpy
P
ancestor
x
418.1X#1
F1
F2
Selection: Dpy and WT
Selfing
(4-5 times)
2X lines
(x 17 lines)
2X back-cross
Scoring P3.p
(x 3 lines)
Higher frequency of P3.p division
Dpy
P
ancestor
x
418.2X#3.1
F1
F2
Selection: Dpy and WT
Selfing
(4-5 times)
3X lines
(x 6 lines)
3X back-cross
Scoring P3.p
(x 3 lines)
F1*
x
ancestor
F1
F2
Selection: Dpy and WT
Selfing
(4-5 times)
4X lines
(x 15 lines)
4X back-cross
Scoring P3.p
(x 4 lines)

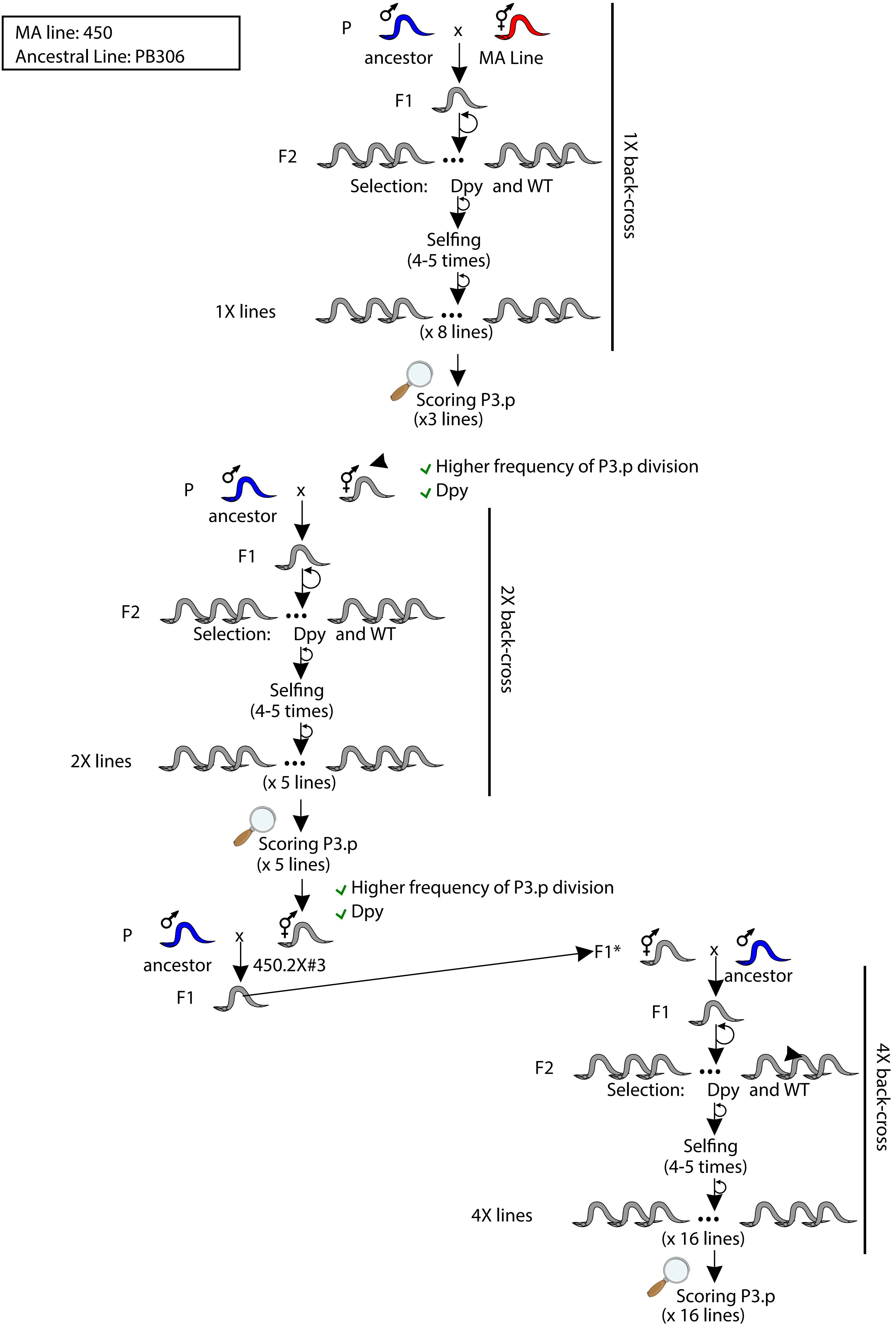

MA line: 450
Ancestral Line: PB306
P
ancestor
x
MA Line
F1
F2
Selection: Dpy and WT
Selfing
(4-5 times)
1X lines
(x 8 lines)
1X back-cross
Scoring P3.p
(x3 lines)
Higher frequency of P3.p division
Dpy
P
ancestor
x
F1
F2
Selection: Dpy and WT
Selfing
(4-5 times)
2X lines
(x 5 lines)
2X back-cross
Scoring P3.p
(x 5 lines)
Higher frequency of P3.p division
Dpy
P
ancestor
x
450.2X#3
F1
F1*
x
ancestor
F1
F2
Selection: Dpy and WT
Selfing
(4-5 times)
4X lines
(x 16 lines)
4X back-cross
Scoring P3.p
(x 16 lines)

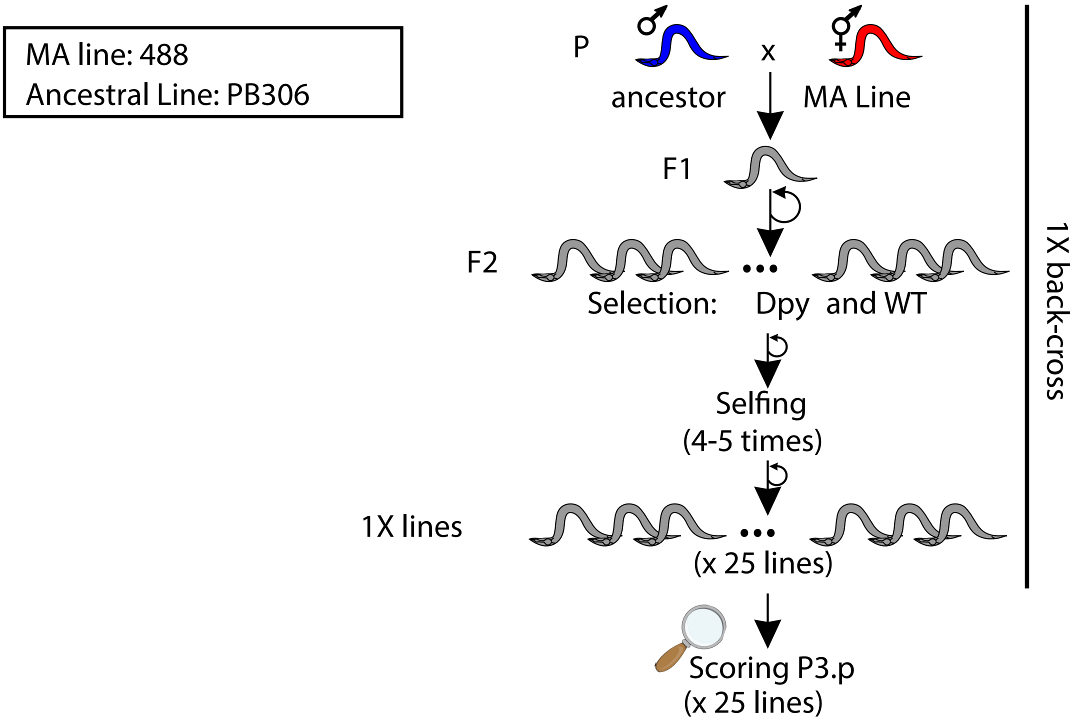
MA line: 488
Ancestral Line: PB306
P
ancestor
x
MA Line
F1
F2
Selection:
Dpy
and WT
Selfing
(4-5 times)
1X lines
(x 25 lines)
Scoring P3.p
(x 25 lines)
1X back-cross

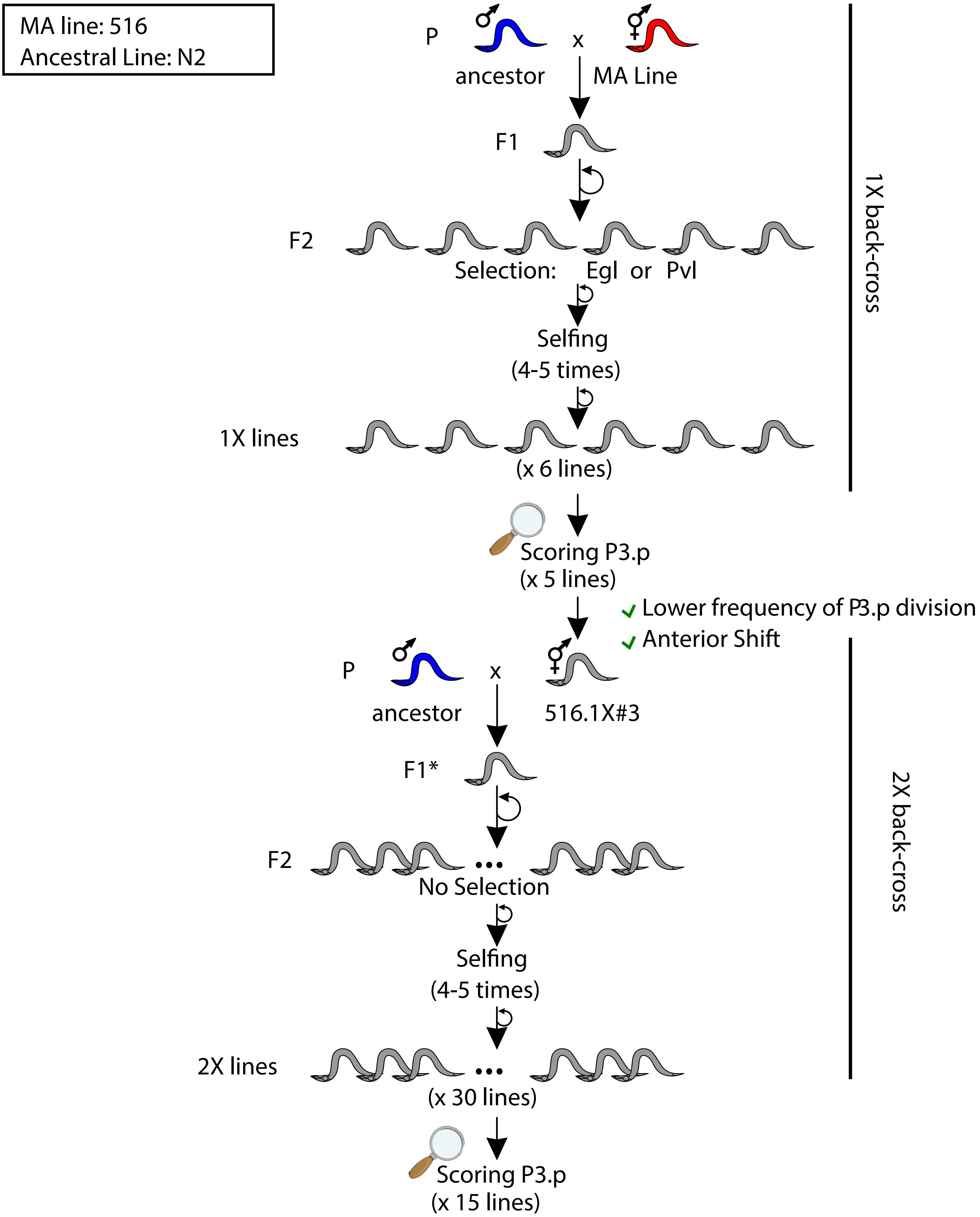
MA line: 516
Ancestral Line: N2
P
ancestor
x
MA Line
F1
F2
Selection: Egl or Pvl
Selfing
(4-5 times)
1X lines
(x 6 lines)
Scoring P3.p
(x 5 lines)
Lower frequency of P3.p division
Anterior Shift
1X back-cross
P
ancestor
x
516.1X#3
F1*
F2
No Selection
Selfing
(4-5 times)
2X lines
(x 30 lines)
Scoring P3.p
(x 15 lines)
2X back-cross

A

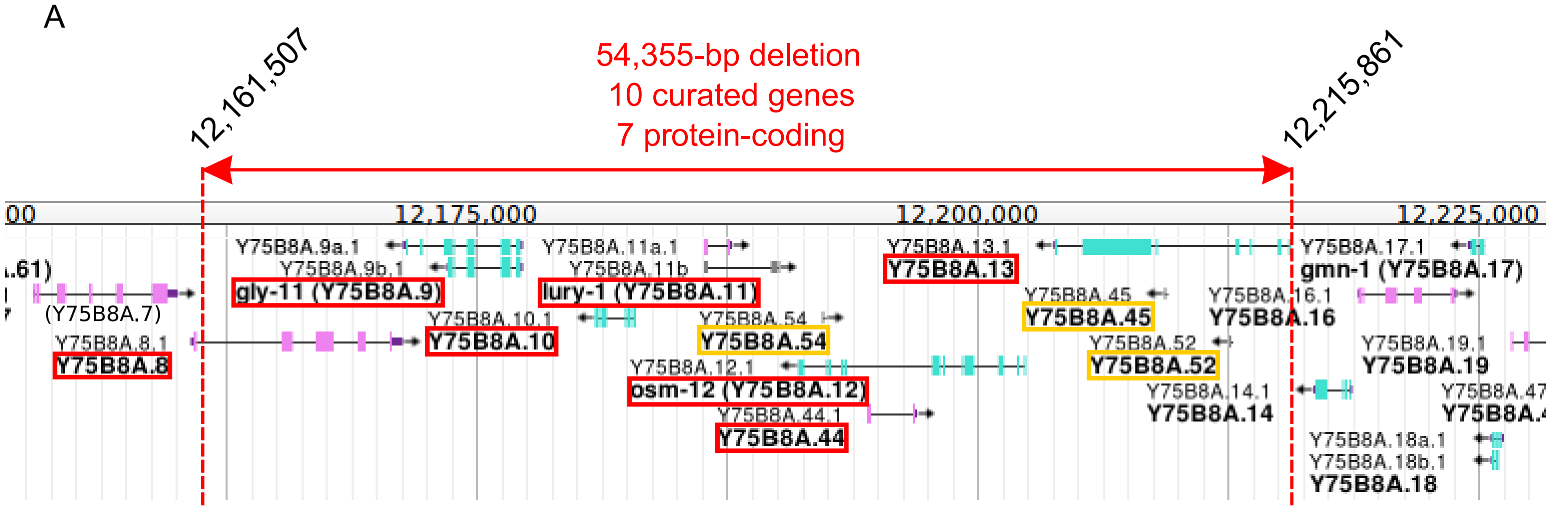


B

| | *Y75B8A.8* | *gly-11 (Y75B8A.9)* | *Y75B8A.10* | *lury-1 (Y75B8A.11)* | *Y75B8A.44* | *osm-12 (Y75B8A.12)* | *Y75B8A.13* |
|---|---|---|---|---|---|---|---|
| Putative null allele | | *gk342* *gk944938* | *tm5064* | *gk961835* *gk961862* | | *n1606* *ok1351* | *gk187847* |
| Targeted deletion | *mf139* | *mf119* *mf120* | *mf135* | | *mf138* | *mf117* | *mf121* |

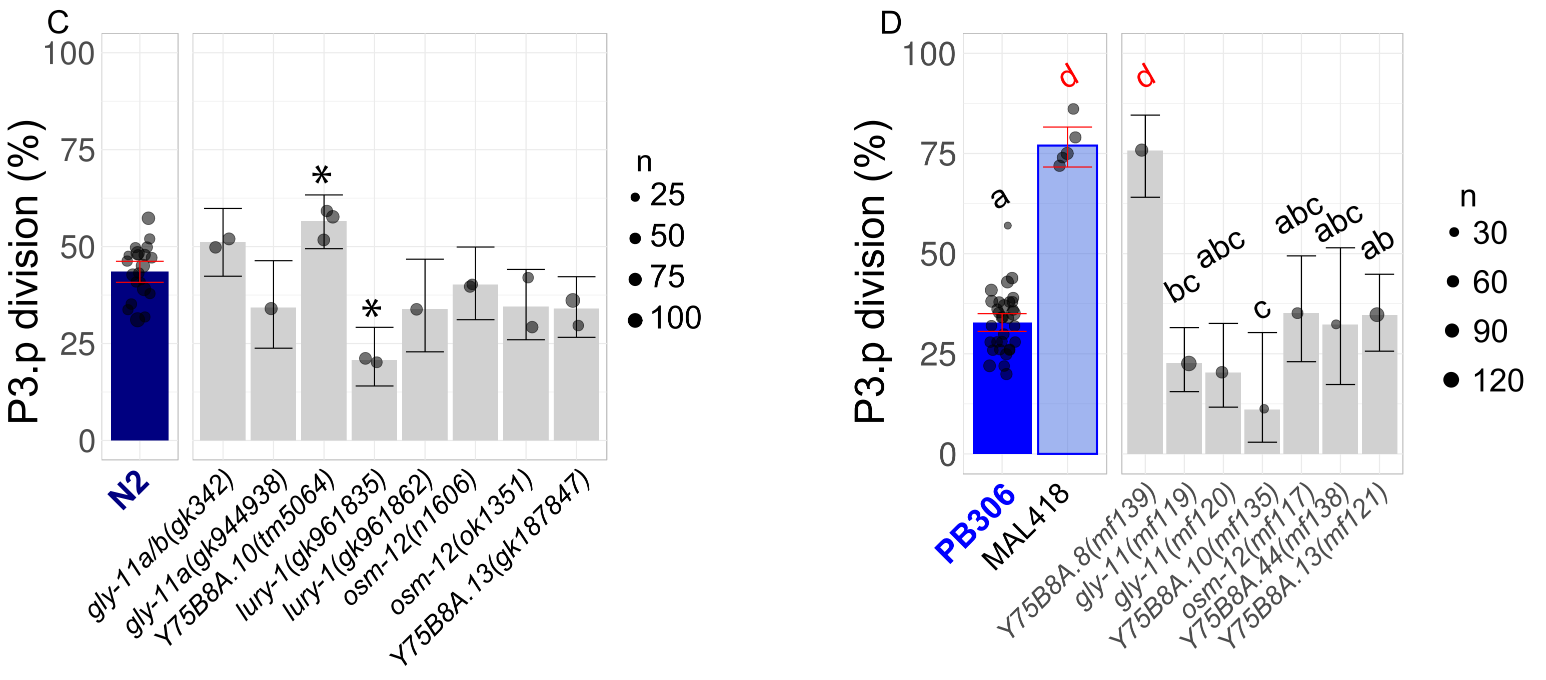


Figure S11

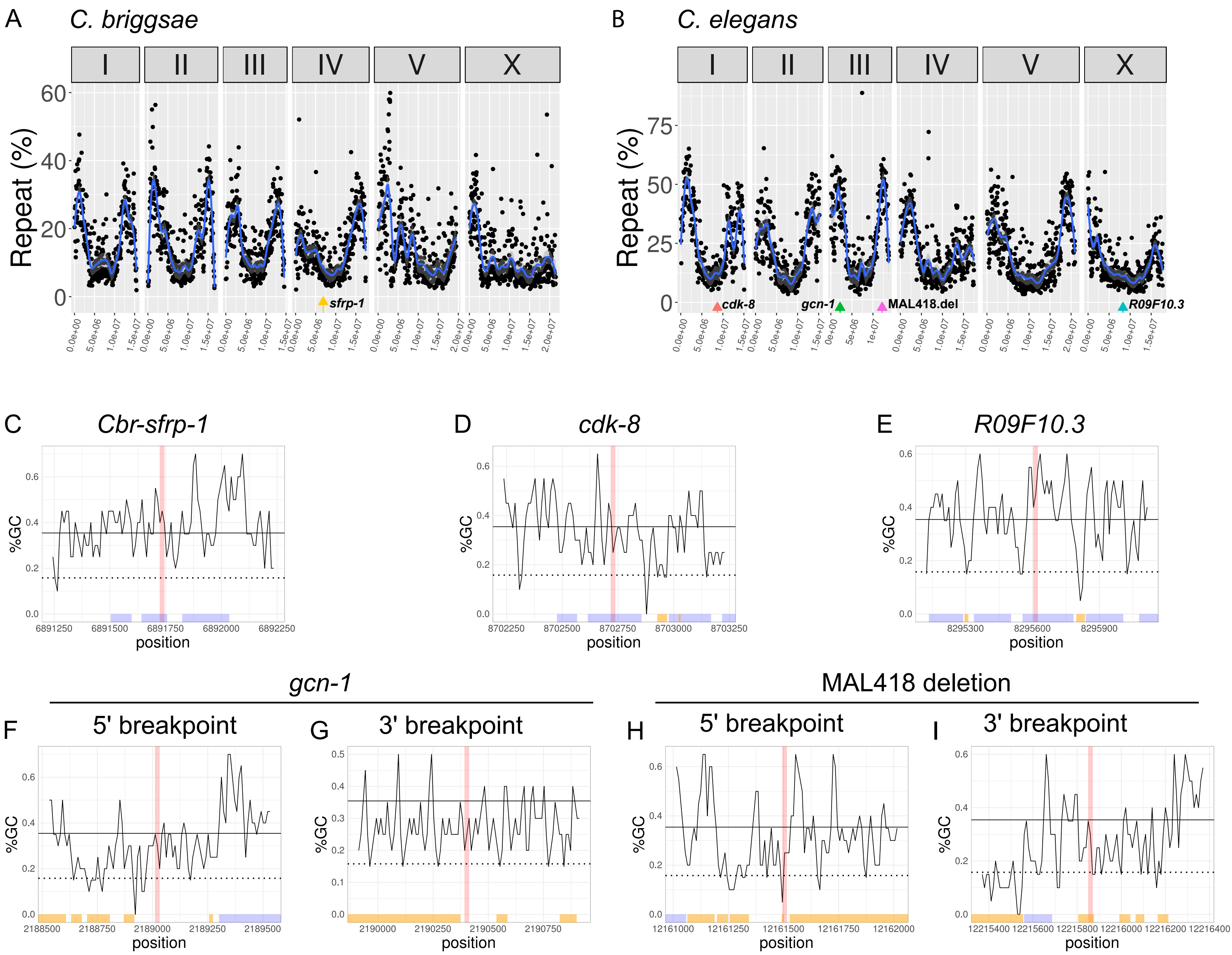
A
C. briggsae
I
II
III
IV
V
X
Repeat (%)
sfrp-1
B
C. elegans
cdk-8
gcn-1
MAL418.del
R09F10.3
C
Cbr-sfrp-1
D
cdk-8
E
R09F10.3
%GC
position
gcn-1
MAL418 deletion
F
5' breakpoint
G
3' breakpoint
H
5' breakpoint
I
3' breakpoint

Figure S12

Mutants related to MAL296 (candidate gene: *sfrp-1* )

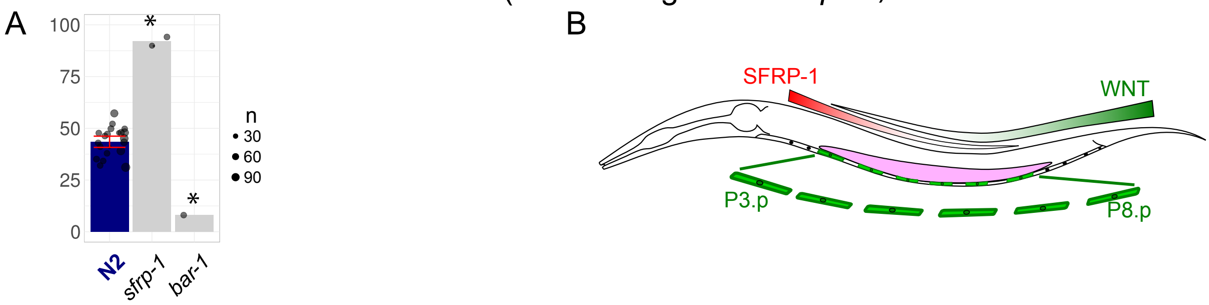


Mutants related to MAL450 (candidate gene: *cdk-8* )

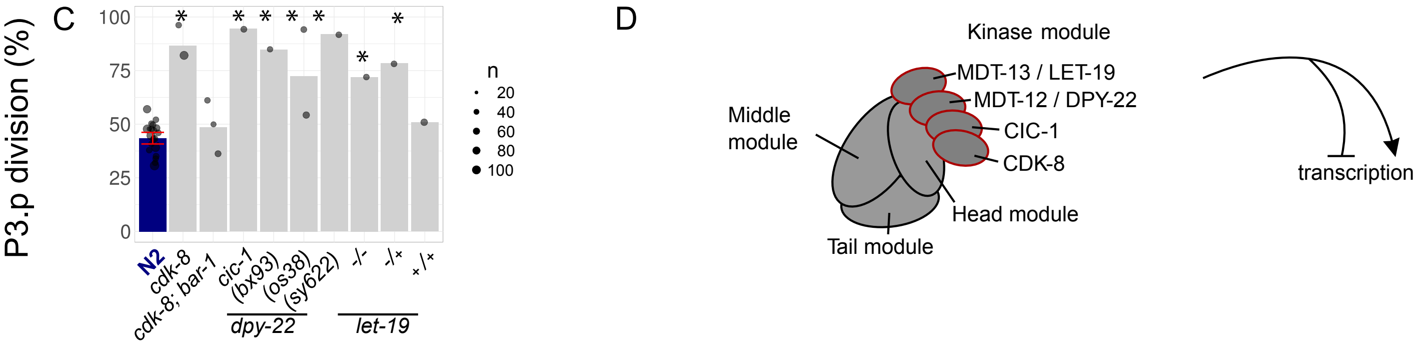


Mutants related to MAL516 (candidate gene: *gcn-1* )

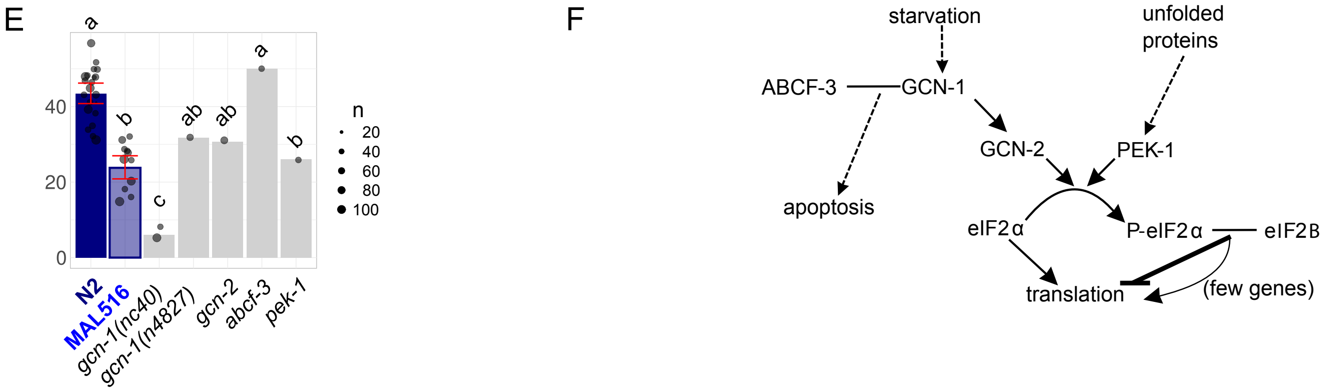


Figure S13
